# Comparative analysis of 2D and 3D distance measurements to study spatial genome organization

**DOI:** 10.1101/076893

**Authors:** Elizabeth H. Finn, Gianluca Pegoraro, Sigal Shachar, Tom Misteli

**Affiliations:** National Cancer Institute, NIH, Bethesda, MD 20892

## Abstract

The spatial organization of eukaryotic genomes is non-random, cell-type specific, and has been linked to cellular function. The investigation of spatial organization has traditionally relied extensively on fluorescence microscopy. The validity of the imaging methods used to probe spatial genome organization often depends on the accuracy and precision of distance measurements. Imaging-based measurements may either use 2 dimensional datasets or 3D datasets including the z-axis information in image stacks. Here we compare the suitability of 2D versus 3D distance measurements in the analysis of various features of spatial genome organization. We find in general good agreement between 2D and 3D analysis with higher convergence of measurements as the interrogated distance increases, especially in flat cells. Overall, 3D distance measurements are more accurate than 2D distances, but are also more prone to noise. In particular, z-stacks are prone to error due to imaging properties such as limited resolution along the z-axis and optical aberrations, and we also find significant deviations from unimodal distance distributions caused by low sampling frequency in z. These deviations can be ameliorated by sampling at much higher frequency in the z-direction. We conclude that 2D distances are preferred for comparative analyses between cells, but 3D distances are preferred when comparing to theoretical models in large samples of cells. In general, 2D distance measurements remain preferable for many applications of analysis of spatial genome organization.

## INTRODUCTION

The eukaryotic genome is functionally organized across several length scales [1,2]. Double-stranded DNA is wrapped around nucleosomes, which are composed of octameric core histones, and further coiled into a chromatin fiber [3], which forms higher order functional conformations, most prominently loops between promoters and enhancers [4,5,6], or between co-regulated genes [7-11]. Furthermore, chromatin forms distinct domains with variable density related to transcriptional activity and histone modifications [12,13]. Large domains of heterochromatin and euchromatin appear to self-associate and also associate with particular nuclear landmarks such as the nuclear lamina [14], the nucleolus [15], and nuclear bodies [16,17]. At the highest level of organization, chromosomes form territories which assume preferred positions within the nucleus [18]. Many of these organizational features have been observed to change during differentiation [19], to be associated with changes in transcription [20,21], and to be disordered in disease [22,23], suggesting that the physical distance between specific genomic locations or relative to nuclear landmarks is an important regulatory feature.

Given their potential regulatory function, measurements of physical distances are of considerable interest. While recently developed biochemical C-methods detect physical interactions between genome regions globally and with relatively high resolution [24], they do not provide spatial distance measurements and can only provide information on pair-wise interactions. The most commonly used means to measure physical distances in the genome is by microscopy-based imaging techniques, particularly fluorescence in-situ hybridization (FISH; [2]).

Measurements of spatial genomic distances can either be performed in 2D or in 3D from a z-stack of images. For most 2D distance measurements, single images for analysis are generated either by maximal projection of z-stacks or by selection of a representative slice. On the one hand, this often simplifies image processing steps and reduces the computational resources needed, especially when high-throughput, automated image analysis is used on large image datasets. On the other, while all the signals in the image are captured, vertical distance information is lost in the projection process. As a consequence, objects spatially separated in 3D space may be detected as co-localizing in the 2D projection if aligned closely along the z- optical axis (Fig. 1A).

**Figure 1.**
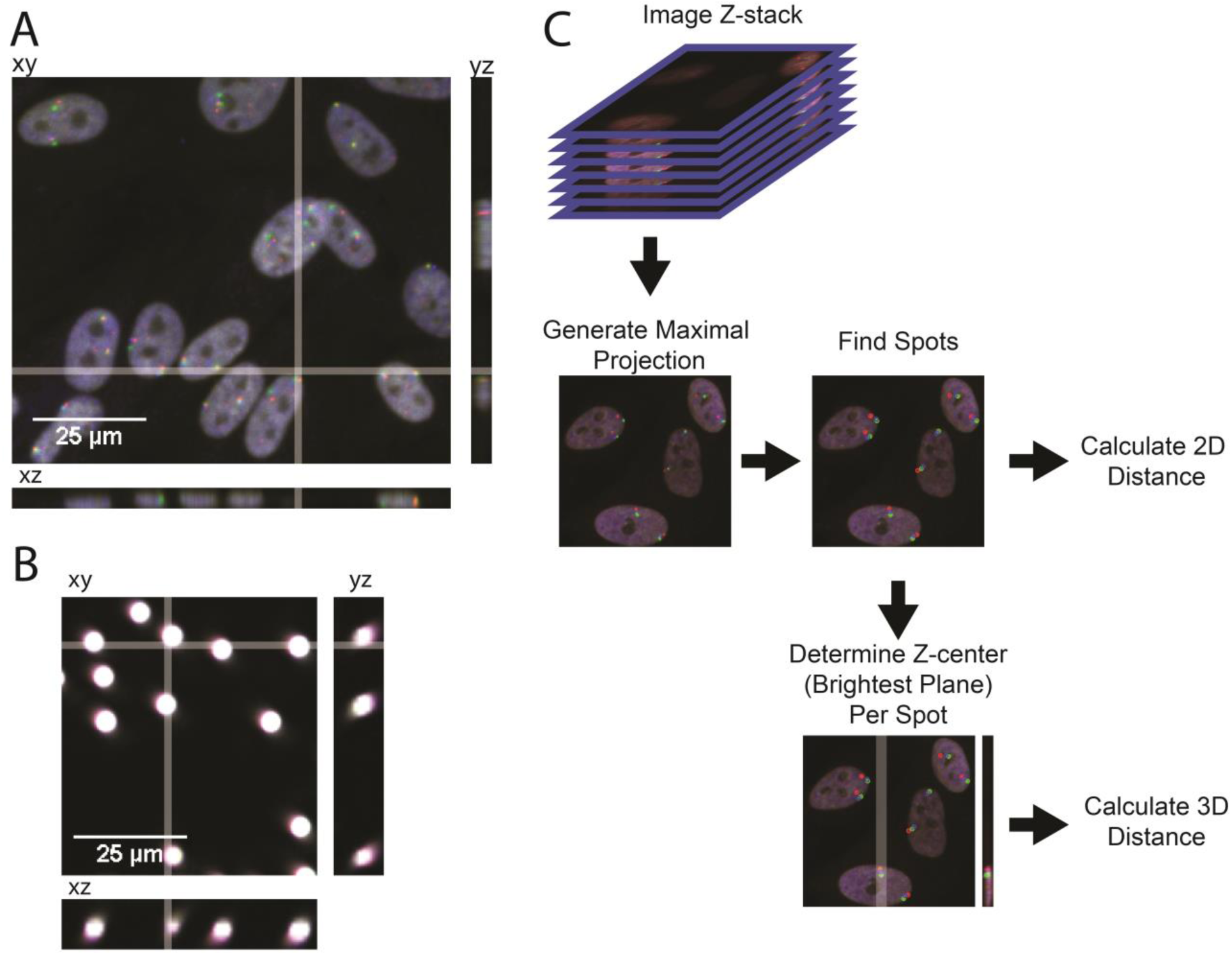
2D and 3D measurements of z-stacks. A: Three orthogonal slices through a field of cells stained for three regions on chromosome 1 (xy, yz, and xz as labeled; xy position of relevant Z-stack marked with white bar). B: Three orthogonal slices through a field of 2.5 m diameter fluorescent beads (xy, yz, and xz as labeled; xy position marked with white bar). C: Example pipeline for generating 2D and 3D distances from a z-stack.

Alternatively, distances between signals can be measured in three dimensions. While 3D datasets preserve all the information contained in the stack, they can introduce artifacts due to low resolution (depth of field), diffraction of signals, and collection of out-of-focus light. Axial resolution is also sensitive to spherical aberrations in lenses, which elongate z-signals ([25,26], Fig. 1A,B), and by chromatic aberrations, which may lead to signal shifts between multiple channels [26]. In addition, when using confocal microscopes, axial resolution depends on the size of the pinhole [27, 28], with a smaller pinhole increasing axial resolution but also decreasing signal, which may lead to signal loss. Finally, variation in the refractive index between cells and media, or between different organelles of the cell, including the nucleus, cause considerable axial distortion in confocal images [27].

Given the advantages and disadvantages of 2D and 3D measurements, we sought to empirically compare the two measurement modalities in different experimental contexts using a spinning-disk confocal microscope, in order to determine the most suitable measurement approach.

## MATERIAL AND METHODS

### Cell Culture

Human foreskin fibroblasts immortalized with hTert (neomycin resistance; [29]) were grown in DMEM media with 10% FBS, 2mM glutamine, and penicillin/streptomycin, and split 1:4 every 3-4 days. Cells were plated in 384 well plates (CellCarrier Ultra, PerkinElmer) at a density of approximately 5000 cells/well between passages 40 and 45 and grown overnight. Plates of cells were then fixed for 10 minutes in 4% paraformaldehyde, washed, and stored in 70% ethanol at −20°C.

### HiFISH Imaging

High-throughput fluorescence in-situ hybridization (HiFISH) was performed as described previously [22,30,31]. Probes were generated via nick translation as described previously [32] from bacterial artificial chromosomes (BACs) to several regions across chromosome 1 (see Table 1). Mixes, reagents, and conditions are exactly as in Meaburn [32], except fluorescently labelled dUTP was substituted for biotin- or digoxigenin- labelled dUTP (Green: ChromaTide Alexa Fluor 488-5-dUTP, ThermoFisher; Red: ChromaTide Alexa Fluor 568-dUTP, ThermoFisher; Far-red: Amersham CyDye Cy5-dUTP, GE Healthcare) and nucleotide mixes were used at a 1/3 dilution. Probes were mixed, precipitated, and resuspended at a final concentration of 6.67ng/µL in hybridization buffer (50% formamide, 10% dextran sulfate, 1% tween-20, 2X SSC).

**Table 1:** Probes used

| Label | Chr | Start | End | BAC ID |
| --- | --- | --- | --- | --- |
| 11 | 1 | 2,435,023 | 2,569,510 | RP11-1012C20 |
| 52 | 1 | 12,768,721 | 12,925,598 | RP11-380A17 |
| 74 | 1 | 18,566,410 | 18,766,881 | RP11-164D21 |
| 80 | 1 | 20,012,370 | 20,177,633 | RP11-451I3 |
| 88 | 1 | 22,000,503 | 22,169,684 | RP11-655O19 |
| 91 | 1 | 22,549,855 | 22,721,150 | RP11-702L8 |
| 354 | 1 | 88,222,723 | 88,418,435 | RP11-647L1 |
| 360 | 1 | 89,759,076 | 89,913,487 | RP11-465E12 |
| 375 | 1 | 93,528,778 | 93,687,931 | RP11-657I7 |
| 385 | 1 | 96,067,993 | 96,226,767 | RP11-489I11 |
| 389 | 1 | 97,025,756 | 97,207,861 | RP11-357E20 |
| 422 | 1 | 105,256,971 | 105,427,281 | RP11-75M16 |
| 463 | 1 | 115,549,942 | 115,729,562 | RP11-675C19 |
| 476 | 1 | 118,815,747 | 118,989,849 | RP11-419G7 |

Fixed cells were permeabilized for 20 min in 0.5% saponin/0.5% Triton X-100/PBS, washed twice in PBS, incubated for 15 min in 0.1N HCl, neutralized for 5 min in 2X SSC, and equilibrated for at least 30 min in 50% formamide/2X SSC before probes were added. Probes and nuclear DNA were denatured at 85°C for 7.5 min and plates were immediately moved to a 37°C humid chamber for hybridization overnight. The next day, plates were washed thrice in 1X SSC, thrice in 0.1X SSC, stained with DAPI, mounted in PBS, and imaged.

Imaging was performed in four channels (405, 488, 561, 640 nm excitation lasers) in an automated fashion using a spinning disk high-throughput confocal microscope (PerkinElmer Opera QEHS) using a 40x water immersion lens (NA = 0.9) and pixel binning of 2 (pixel size = 320 nm). Approximately 1,000 cells in 20-40 fields were imaged per well. For sparse sampling, z-stacks of 7 m thickness with images spaced 1µm apart were acquired in three separate exposures. For Nyquist sampling, z-stacks with total 4.2 m thickness and 300 nm image intervals were generated. In all exposures the light path included a primary excitation dichroic (405/488/561/640 nm), a 1^st^ emission dichroic longpass mirror: 650/ 660- 780, HR 400-640 nm and a secondary emission dichroic shortpass mirror: 568/ HT 400- 550, HR 620-790 nm. In exposure 1, samples were excited with the 405 and 640 nm lasers, and the emitted signal was detected by two separate 1.3 Mp CCD cameras (Detection filters: bandpass 450/50 nm and 690/70 nm, respectively). In exposure 2, samples were excited with the 488 nm laser and the emitted light was detected through a 1.3 Mp CCD camera (Detection filter: bandpass 520/35). In exposure 3, samples were excited with the 561 nm laser and the emitted light was detected through a 1.3 Mp CCD camera (Detection filter: bandpass 600/40).

### 2D and 3D Image Analysis

Automated analysis of all images was performed based on a modified version of a previously described Acapella 2.6 (PerkinElmer) custom script [30, 33-35]. This custom script performed automated nucleus detection based on the maximal projection of the DAPI image (ex. 405 nm) to identify cells. Spots within these cells were subsequently identified in maximal projections of the Green (ex 488 nm), Red (ex. 561 nm) and Far Red (ex. 640 nm) images, using local (relative to the surrounding pixels) and global (relative to the entire nucleus) contrast filters. The x and y coordinates of the central pixel in each spot were calculated. The z coordinate of the spot center was then calculated by identifying the slice in the z-stack with the highest value in fluorescence intensity for each of the spot centers. Datasets containing x,y and z coordinates for spots in the Green, Red and Far Red channels as well as experiment, row, column, field, cell, and spot indices, were exported from Acapella as tab separated tabular text files. These coordinates datasets were imported in R [36]. 2D and 3D distances for each pair of Red:Green, Red:Far Red, or Green:Far Red probes within a cell were generated on a per-spot basis using the SpatialTools R package [37]. Subsequent analyses were performed in R using the plyr [38], dplyr [39], ggplot2 [40], data.table [41], knitr [42] and stringr [43] packages. All images, scripts, and datasets are available upon request.

### Statistical Analysis

For 2D/3D scatterplots, 2D and 3D distances were calculated on a per-green-spot basis using the SpatialTools R package [37]. These distances were plotted using ggplot2 [40].

For modelled 2D/3D scatterplots, random pairs of coordinates were generated with a normal distribution; standard deviation was 100 for x and y and 10, 30, 50, or 100 for z. Distances between these pairs were calculated in 2D and 3D using the SpatialTools R package [37] and these points were plotted using ggplot2 [40].

For 2D/3D colocalization frequencies in fibroblasts, both 2D and 3D distances were calculated using the SpatialTools R package [37]. Minimal distances were calculated on a per-green-spot basis using the data.table R package [41]. The percentage of spot pairs within thresholds of 350 nm, 700 nm, 1 µm, 2 µm, 3 µm, and 4 µm was calculated using the data.table R package [41] and plotted using ggplot2 [40].

For triplet associations in fibroblasts, both 2D and 3D minimal distances were calculated on a per-green-spot basis as above. Triplets were defined as events where a single green spot was within 1 µm of a red spot and a far-red spot. Both pairing frequencies within 1µm, and triplets, were counted. For each probe set, expected proportions of triplets were calculated as p(G:R)*p(G:F) where p(G:R) is the proportion of green and red spots colocalizing within 1µm and p(G:F) is the proportion of green and far red spots colocalizing within 1µm. 95% confidence intervals were calculated with the modified Wald method.

For distance distribution histograms in fibroblasts, minimal distances were calculated on a per-green-spot basis and plotted.

## RESULTS

### 2D vs 3D distance measurement over varying length scales

In order to determine how well 2D and 3D distance measurements correspond to each other, we systematically compared distance measurements across several length scales using 2D or 3D measurement regimes. As a model system we used a series of probes tiling two regions of approximately 20 Mb each on chromosome 1 (diagrammed in Fig. S1). Each probe pair was separated by at least 10 Mb and in total 20 different probe pairs were examined. We determined pairwise distances between each pair of loci, both in 2D from maximal projections and in 3D using automated high-throughput spinning disc confocal microscopy as described in figure 1C (Materials and Methods). We then compared 2D and 3D distances for each pair of spots in each cell by pooling in silico all the probe pairs studied (Fig. 2A). We observe a strong distance dependence for the concordance of 2D and 3D distances. 2D and 3D distances between spot centers are in good agreement for distances above approximately 5 µm with average discrepancy between measurements of less than 1% (Fig. 2A). Below 5 µm we see on average a 29% difference between 2D and 3D distances, and below 1 µm we see on average an 83% difference. Similarly, while we observe a statistically significant difference between 2D and 3D distances overall (two sample t-test, p=7.057e-13), this difference is much less significant for points separated by at least 5 µm (two sample t-test, p=0.01744).

**Figure 2.**
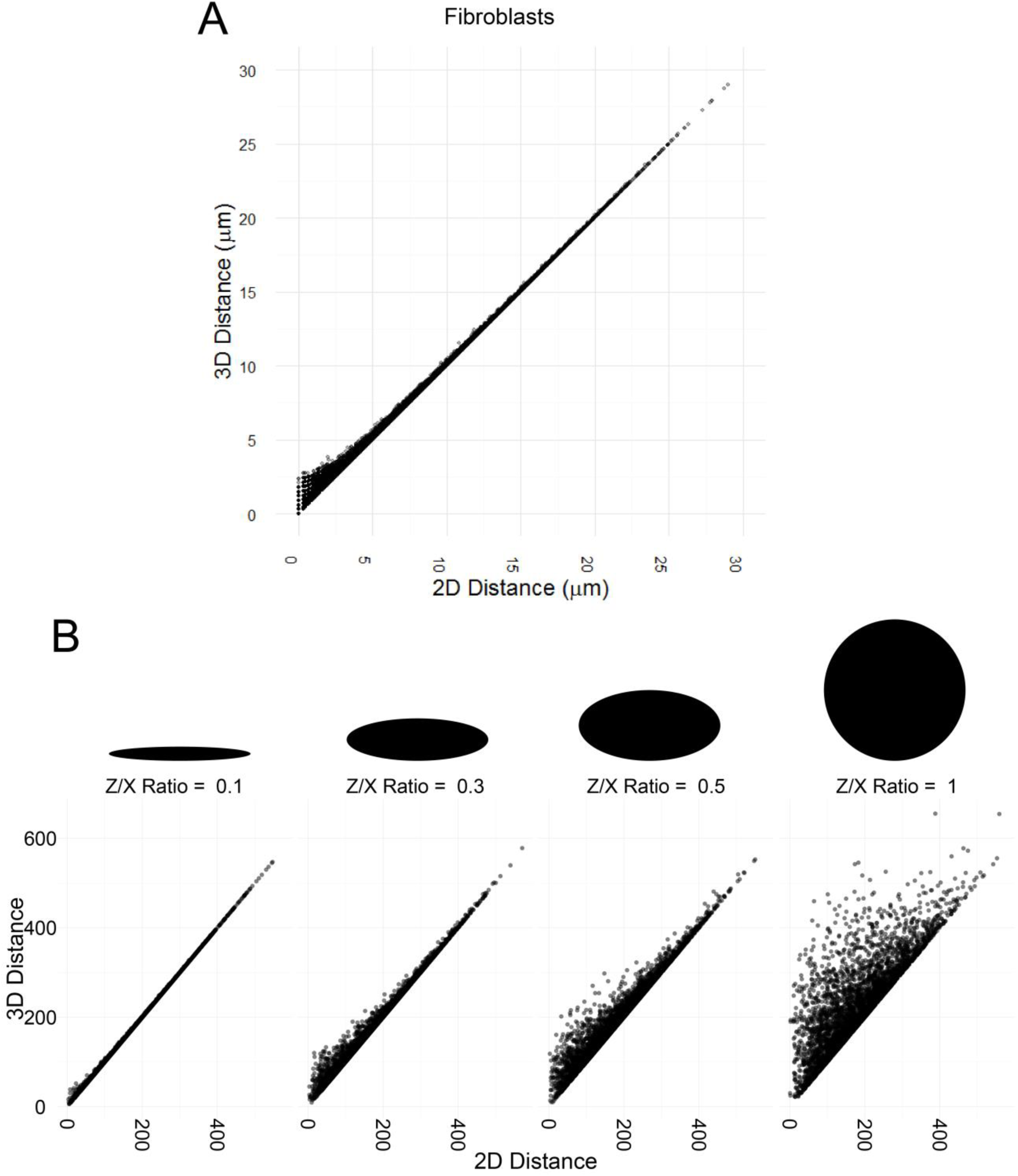
2D vs 3D distances in nuclei of multiple shapes. A: Scatterplot showing 2D vs 3D distance for minimal distances between FISH signals in fibroblasts. B: Scatterplots showing modeled relationship between 2D and 3D distances at random points in volumes of various sizes.

Prompted by the fact that the 5 µm cut-off is within the dimensions of the average height of the fibroblast cell nucleus, we interrogated the effect of nuclear shape on 2D and 3D distance measurements by generating theoretical models for 2D and 3D distance distributions in nuclei with varying degrees of flatness (Fig. 2B-E). In our simulated models, the x and y coordinates were sampled from a random normal distribution with a mean of 300 arbitrary units and a standard deviation of 100 arbitrary units; the z coordinate was set with a mean of 300 and standard deviations of 10 (z/x = 0.1), 30 (z/x = 0.3), 50 (z/x = 0.5) and 100 (z/x = 1). In scatter plots of 2D versus 3D distance for all modeled points, we observe that flatter nuclei show more agreement between 2D and 3D analysis, and that agreement is stronger at larger distances. Consistent with our observations in fibroblasts, deviations are greater at shorter distances, and in addition we observe a striking effect of nuclear shape: considering all spots, average discrepancy between measurements was 21% for a model of perfectly round nuclei (z/x = 1), 10% for z/x = 0.5, 6% for z/x = 0.3, and only 1% for our flattest model (z/x = 0.1). Considering only spot pairs within 75 units, which is 15% of the maximum radius, or roughly 5 µm in a fibroblast with maximum distance between points around 30 µm, average discrepancy between measurements is still 21% for the round model, but increases to 21% for z/x = 0.5, 17% for z/x = 0.3, and 5% for the flattest model. Similarly, the statistically significant difference between 2D and 3D distances is observable mostly in round cells and at short distances. When considering all pairs, we observe highly significant differences between 2D and 3D modeled differences for z/x = 1 (p < 2.2e-16), and z/x = 0.5 (p = 1.379e-13), significant differences when z/x = 0.3 (p = 0.001228), and no significant differences when z/x = 0.1 (p = 0.7132, two sample t-test). Considering only pairs separated by at least 75 units, we see a general decrease in significance (z/x = 1: p < 2.2e-16, z/x = 0.5: 2.067e-10, z/x = 0.3: 0.009486, z/x = 0.1: p = 0.7753). It is worth noting here as well that the difference in the means for z/x = 0.3 with point pairs separated by at least 75 units is only 5 units (2D mean: 197.0418, 3D mean: 202.2165) which corresponds to approximately 333 nm, which may be within normal noise due to drift of the microscope and aberrations in mirrors and lenses. Thus, both cell shape and average distance between spots affect the ratio of measurements made in 2D to those made in 3D. Overall, at longer distances and in flatter cells, 2D and 3D measurements are not significantly different.

### 2D and 3D association probabilities

Since 2D and 3D distances diverge at short ranges, it seems likely that association probabilities between genome regions in 3D space, which rely on detection of short distance measurements, would differ depending on whether they are calculated based on 2D or 3D distances. We tested this hypothesis by comparing 2D and 3D co-localization frequencies for pairs of regions on chromosome 1, separated by between 2 and 20 Mb (Fig. S1A). For analysis, we considered varying association thresholds between 350 nm and 4 µm (Fig. 3). We observe that the distance threshold has a notable effect on the frequencies calculated from 2D or 3D distances. In particular, deviations were more noticeable at shorter thresholds. With a threshold of 3 or 4 µm, we see little difference between 2D and 3D frequencies (Fig. 3A, green and yellow dots) whereas with a threshold of 1 µm or less, deviations are greater (Fig. 3A, orange, dark blue, and gray dots). Furthermore, we observe greater effects at loci which interact more frequently; for example, two loci which colocalize in 25% of cells in 3D appear to interact in as many as 55% of cells in 2D, whereas two loci which colocalize in under 10% of cells in 3D colocalize in under 20% of cells in 2D (Fig. 3A). These observations suggest that 2D and 3D analysis of rare associations and long-range interactions are equivalent, however, frequent, short-range associations may be overestimated in 2D measurements. Hence, for the study of close physical associations between loci, 3D distance measurements are likely to be more accurate.

### Accuracy of 2D vs 3D distance measurements

The accuracy of a measurement is defined as closeness of agreement between the measurement and the true value of the measured entity. The higher accuracy of 3D measurements should lead to an improved ability to detect deviations from expected relationships. One very simple such theoretical model is that two pairwise interactions between one bait and two targets are independent. That is to say, that a co-localization between locus A and locus B does not change the probability of a co-localization between locus A and locus C. With this assumption, we are able to make a prediction of the number of triplets (A, B, C) that we observe. However, it is likely that biological factors, such as chromatin state, transcriptional activity, and protein binding play a role in both of the pairwise interactions, and that in some cases the triplets will occur more often, or less often, than expected. If 3D measurements are more accurate than 2D measurements, they will be more sensitive to deviations from a neutral model. To test this hypothesis, we used 2D and 3D measurements to examine clustering behavior at four triplets spanning 20Mb on chromosome 1 (from 2,301,890 to 22,549,855).

**Figure 3:**
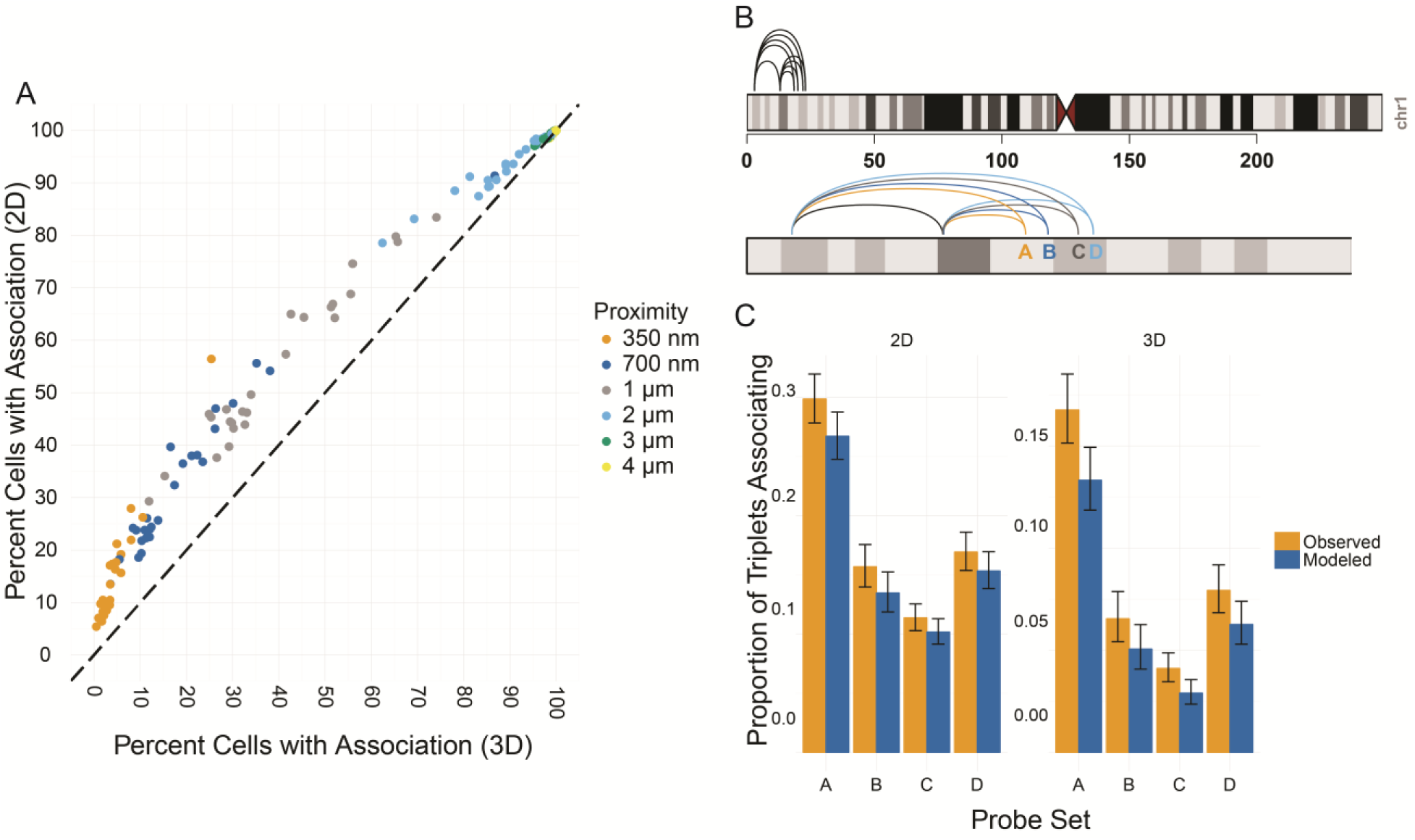
Co-localization proportions in 2D and 3D. A: Scatterplot showing co-localization proportion based on 2D distances vs. co-localization proportion based on 3D distances. Distance cutoff color coded. B: Ideogram showing probe assignment into triplets. C: Bar graph showing proportion of green spots interacting with red and far-red spots simultaneously. Orange: observed; blue: expected.

Triplets were chosen from among our pairwise associations (Fig. 3B). Each probe triplet contained our most frequently occurring interaction, consisting of a bait between 12.75 and 13 Mb on chromosome 1 and a target 10 Mb upstream. The third probes mapped to a region approximately 10 Mb downstream from the bait and were spaced by 0.5-2 Mb. We modeled triplet associations, assuming independence between each pairwise association, and compared observed frequencies of clustered interactions detected by 2D or 3D analysis to modeled frequencies. We observe a statistically significant tendency for probes to cluster more than expected in both 2D and 3D (Fig. 3C). The enrichment and significance were greatly enhanced in 3D. The interaction frequency in 2D is on average 1.12x that of the expected value (p-value: 5.133E-5), in 3D the interaction frequency is on average 1.29x that of the expected value (p-value: 1.329E-10, two-tailed proportion t-test). Finally, while we see differences between observed rates of triplet associations and those modeled based on pairwise association frequencies at all four triplets in 3D, we find individual triplets were rarely significantly enriched (p-value: 0.01) in 2D (Fig. 3C). Thus, the inaccuracy inherent in 2D distance metrics makes it difficult to detect statistically significant trends in the data, whereas the improved accuracy with 3D distance thresholds identifies such interactions.

### Variability and noise in 2D vs 3D datasets

When examining overall distance distributions in 2D and 3D, we noticed periodic noise in 3D distances taken with a 1 µm z-stack which was absent in 2D (Fig. 4A middle column). This noise could plausibly come from poor z-resolution. To determine whether noise caused by decreased resolution in Z significantly deforms distance distributions, we compared distance distributions for 2D distance measurements and 3D distance measurements generated from z-stacks with a 1 µm slice or a 300 nm slice (Fig. 4A). 3D distance measurements generated from z-stacks with a 1 µm slice contained prominent periodicity in distance distributions which were absent in 2D distance measurements and more densely sampled 3D distance measurements (Fig. 4A). The discontinuities in 3D distances were most visible at 1 and 2 µm distances, but were visible in distances up to 4 µm. The 3D distance measurements have a period of approximately 1 µm, equivalent to the spacing between z-sections in our dataset. It is worth noting that our x-y distances also represent discrete measurements with an x-y pixel size of 320 nm. As such, we observe discontinuities in all distance measurements at very short length scales (<1µm) (Fig. 4B), however, when the resolution in the x-y is greater than in z, discontinuities at very low distances in the x-y plane do not significantly alter the overall distance distributions. We conclude that 3D distances are sensitive to noise generated by decreased resolution across the optical axis.

**Figure 4:**
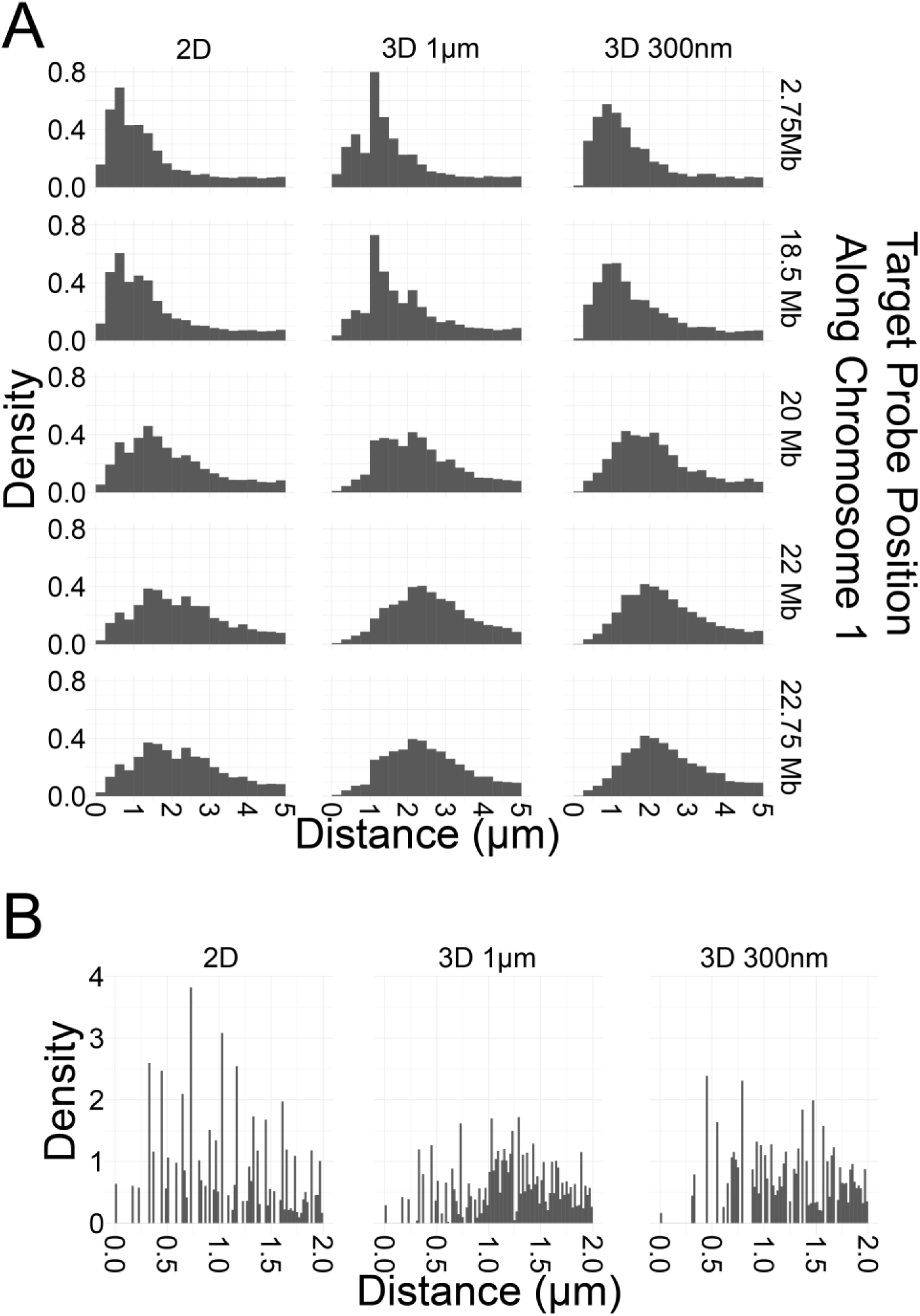
Various observed artifacts in 3D. A: Histograms of distance distributions for minimal distances in fibroblasts for various probe sets with various average distances. Discontinuities due to lowered resolution in z are visible in 3D distance distributions with 1 µm z-slices but not 2D distance distributions or 3D distance distributions with 300 nm z-slices. B: Discontinuities due to resolution in x-y are visible at very short distances with very small binning regardless of the method for calculating distances.

## DISCUSSION

Determination of distances in the cell nucleus is routinely performed in 2D, mostly due to the ease of measurements and the ability to analyze a larger number of individual cells. The use of 2D measurements has become standard, despite the fact that, intuitively, measurements of distance in 3D should be more accurate than in 2D. However, technical and physical limitations of microscopes mean that the vertical position of a signal is difficult to pinpoint exactly, and as such measurements in 3D may be overall less precise than measurements in 2D. We have here systematically compared 2D vs 3D distance measurements using FISH datasets.

We do not find that 3D distance measurements are always better than 2D measurements, suggesting that use of 2D analysis is a valid approach to studying nuclear organization. In fact, for some cases, such as determination of large distances, particularly in flat cells, 2D and 3D distances will yield very similar results. Furthermore, when the computational requirements for 3D cells become onerous, when the need for precision is high, or when only a few cells can be sampled, 2D measurements are advantageous. In addition, calculating 2D distances using only a single slice may speed acquisition time, reduce photobleaching and phototoxicity, and facilitate applications such as live cell microscopy. In addition, most practical applications are comparative in nature: the distances or association frequencies between a test pair of loci are compared with a set of control loci analyzed the same way. Provided that the measured distances for the test and the control pairs are similarly distributed they should be affected by the same systematic errors in measurements, thus making an internal comparison valid, regardless of whether 2D or 3D measurements are used.

When considering measurement of short-range interactions, the choice between 2D and 3D distances is determined by the need of accuracy, defined as the reflection of the measurement of the ground truth, versus precision, defined as the internal consistency of multiple independent measurements, versus the time taken to acquire images (Table 2). 2D distance measurements can be generated from a single projected image, and as such are favored when rapid computation is required, including in high-throughput experiments involving tens of thousands of fields of view. They are also precise by nature, due to better resolution in x and y, and easier normalization for optical aberrations in the x-y plane. 3D distances are more accurate, however, in order to gain comparable precision in 3D distance calculations to that achieved in 2D calculations, it is necessary to use relatively thin z-slices (300 nm z-slices in our case) rather than sampling at the depth of field of the objective lens used (1 µm z-slices in our case). In particular, the optimal height for a z-slice can be determined analytically based on the objective and the wavelength of the fluorophores used, according to the Nyquist-Shannon sampling theorem, which determines the minimal sampling density needed to capture all information from the microscope into the image. One rule of thumb is to sample at one half the Rayleigh criterion, r = 0.61 λ/NA for wavelength λ and numerical aperture NA (for our experimental set up this yields a suggested sampling rate between 270 and 440 nm). Another suggestion, which addresses the fact that z-resolution is often limited by pinhole size and depth of field rather than diffraction of light, is to sample at one half or one third the maximal resolution (for our experimental set up with 1 µm depth of field this yields a suggested sampling rate between 300 and 500 nm). Unfortunately, this greatly increases the number of exposures required for imaging and thereby greatly slows acquisition time. The ‘ground truth’ is most accurately reflected by 3D distances generated from image stacks with high z-slice resolution. However, for applications where many conditions must be sampled, and therefore acquisition time is at a premium, this approach is prohibitive. In these cases, 2D distances or 3D distances generated from sparser z-stacks are preferred. When precision is crucial, for instance when examining the correlation between two parameters, 2D distances may be preferred. On the other hand, when accuracy is crucial, for instance when comparing to a theoretical model, sparse 3D distances are preferred. Thus, given the generally good alignment of 2D and 3D measurements, the choice between 2D and 3D distance measurements for frequent interactions depends on the biological and statistical question asked (Table 2).

**Table 2:** Comparison of imaging modalities

|  | **2D** | **3D**<br>1µm z-stack<br>resolution | **3D**<br>300nm z-stack<br>resolution |
| --- | --- | --- | --- |
| Fast? | Yes | Yes | No |
| Precise? | Yes | No | Yes |
| Accurate? | No | Yes | Yes |

Although not specifically tested here, measurements of radial gene positioning, i.e. the location of a gene relative to the center of the nucleus, likely benefit from 2D analysis compared to 3D measurements [24]. Because radial position is often computed as a percent of the radius of the cell, in each measurement a long distance is being considered. The edge of the nucleus is more computationally difficult to find than a second spot center, and such computations become much more difficult in 3D, especially in a high-throughput format, putting a premium on computational resources. In addition, there are several ways to reduce the bias generated by the maximal projection. Since radial position is most often considered in a comparative fashion, the systematic bias resulting from loci at the ‘top’ or ‘bottom’ of the nucleus appearing in the center in a focal plane or maximal projection will be, essentially, considered in the background measurements. In addition, since radial position is measured on a per-spot basis, it is possible to remove some of the most biased spots by selecting a single central focal plane, ruling out spots at the very top or bottom of the nucleus. Thus, for measurements of radial position it is likely that the advantages of using 3D distances are limited and 2D distance measurements are preferred in most cases.

In sum, we find here, that both 2D and sparse 3D measurements have systematic biases that must be taken into account. In 2D, interactions appear more likely than they are in reality due to the overlay effect created when generating a maximal projection. In sparse 3D analysis, noise is added due to poor z-resolution. To properly use 2D distances, systematic errors must be eliminated either by empirically measuring background values, or by cross-comparing multiple samples which will all have the same bias, rather than using a theoretical model. To properly use 3D distances, it is imperative that a large number of cells is imaged in order to minimize the effect of the additional sources of noise such as chromatic aberrations or imperfections in the imaging light path. Taken together, our comparative analysis shows that both 2D and 3D distance measurements are imperfect, but suggests good agreement between the two approaches. While specific experimental designs may require the strict use of either method, the logistically simpler and practically faster 2D distance measurements will likely be preferred in many situations and will be appropriate in most cases.

## ACKNOWLEDGEMENTS

This research was supported, in part, by the Intramural Research Program of the National Institutes of Health (NIH), National Cancer Institute, and Center for Cancer Research and by the 4D Nucleome Common Fund (5U54DK107980-01).

---

**Figure S1:**
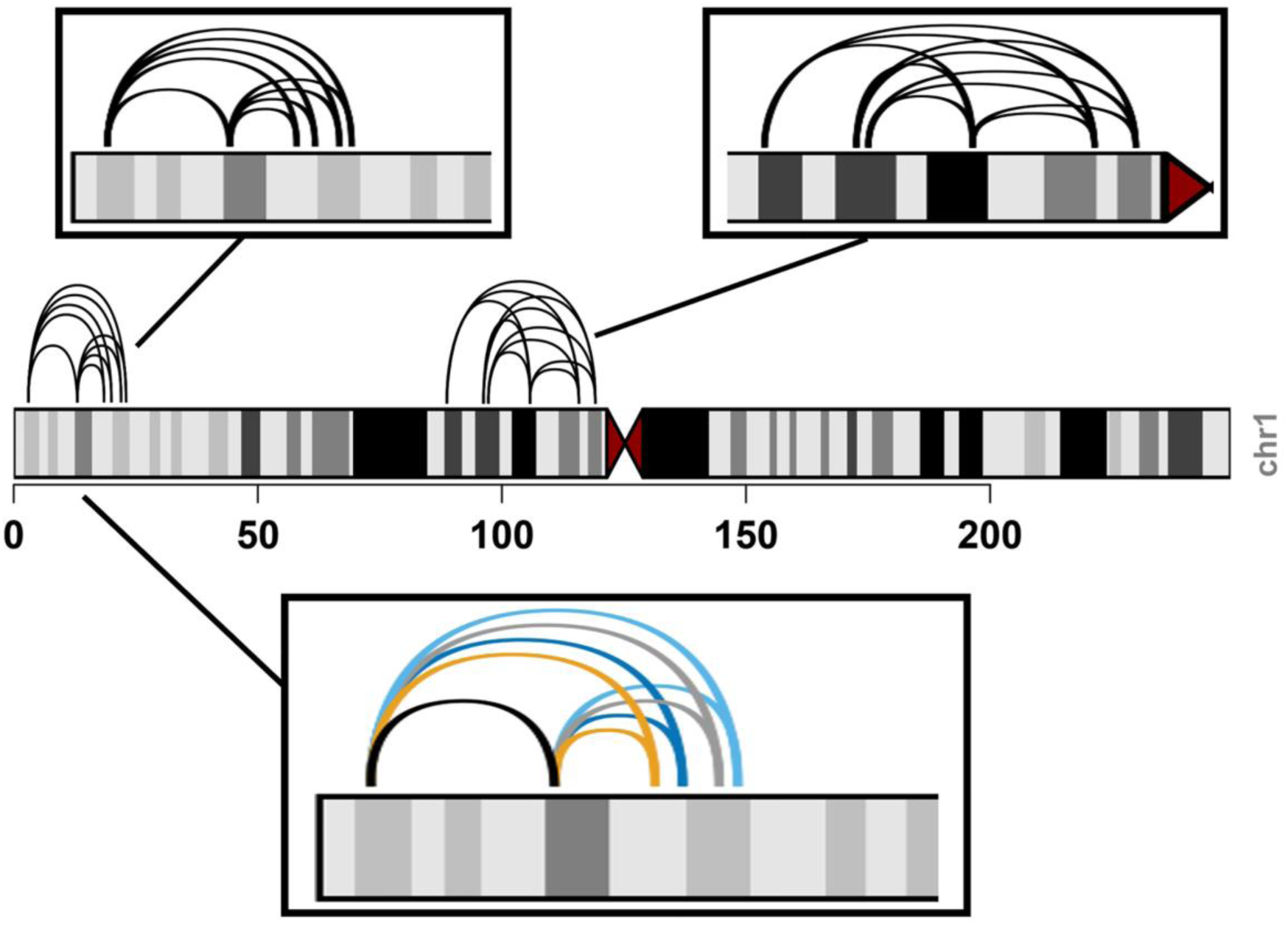
Ideograms showing the placement of probe pairs (arcs) on chromosome 1. A, B: Zoomed-in view of all probe pairs in two tiled 20Mb regions. C: Color-coded view of all triplets in the upstream of two tiled 20Mb regions.

